# CryoTEN: Efficiently Enhancing Cryo-EM Density Maps Using Transformers

**DOI:** 10.1101/2024.09.06.611715

**Authors:** Joel Selvaraj, Liguo Wang, Jianlin Cheng

## Abstract

**Motivation:** Cryogenic Electron Microscopy (cryo-EM) is a core experimental technique used to determine the structure of macromolecules such as proteins. However, the effectiveness of cryo-EM is often hindered by the noise and missing density values in cryo-EM density maps caused by experimental conditions such as low contrast and conformational heterogeneity. Although various global and local map sharpening techniques are widely employed to improve cryo-EM density maps, it is still challenging to efficiently improve their quality for building better protein structures from them.

**Results:** In this study, we introduce CryoTEN - a three-dimensional U-Net style transformer to improve cryo-EM maps effectively. CryoTEN is trained using a diverse set of 1,295 cryo-EM maps as inputs and their corresponding simulated maps generated from known protein structures as targets. An independent test set containing 150 maps is used to evaluate CryoTEN, and the results demonstrate that it can robustly enhance the quality of cryo-EM density maps. In addition, the automatic de novo protein structure modeling shows that the protein structures built from the density maps processed by CryoTEN have substantially better quality than those built from the original maps. Compared to the existing state-of-the-art deep learning methods for enhancing cryo-EM density maps, CryoTEN ranks second in improving the quality of density maps, while running *>* 10 times faster and requiring much less GPU memory than them.

**Availability and implementation:** The source code and data is freely available at https://github.com/jianlin-cheng/cryoten

## Introduction

In a cryogenic Electron Microscopy (cryo-EM) experiment, purified proteins in solutions are fast frozen at cryogenic temperature and then imaged by an electron microscope to obtain their structural information. Compared to traditional techniques (i.e., X-ray Crystallography and Nuclear Magnetic Resonance (NMR)), cryo-EM has the unique capability of determining the atomic structures of large protein complexes and assemblies consisting of multiple protein chains, which are difficult or impossible for other techniques to handle. The reconstruction of protein structures from cryo-EM maps involves three main steps: picking protein particles in 2D cryo-EM micrographs [Dhakal et al., 2024b, Bepler et al., 2020, Gyawali et al., 2024, Dhakal et al., 2024a], aligning the 2D protein particle images from different orientations to reconstruct 3D electron density map [Punjani et al., 2017, Zhong et al., 2021], and using the map for de novo atomic structural model building [Terashi and Kihara, 2018, Terwilliger et al., 2018a, Giri and Cheng, 2024a]. However, one of the main factors affecting the effectiveness of this process is the low contrast and noise present in the 3D electron density map. This problem is partially addressed by various post-processing techniques such as global and local map sharpening that modify the density values of the cryo-EM maps [Terwilliger et al., 2018b, Scheres, 2012, Rosenthal and Henderson, 2003].

Global map sharpening generally involves applying a single B-factor correction to the entire map, aiming to restore contrast and improve interpretability (i.e., the amount of structural information in the map determines the quality of protein structures built from the map). Similar global map sharpening techniques are implemented in phenix.auto sharpen [Terwilliger et al., 2018b], RELION [Scheres, 2012, Rosenthal and Henderson, 2003] post-processing and CryoSPARC [Punjani et al., 2017] sharpening tools. However, assuming a uniform B-factor across the entire map may fail to adapt to the local variations in the map. This could lead to some regions being under-sharpened and other regions being over-sharpened, and cause poor interpretability. Local map sharpening addresses this shortcoming by taking the local variations in cryo-EM map into consideration and adjusting the local regions in the density map accordingly. LocalDeblur [Ramírez-Aportela et al., 2019] uses a wiener filter-based local deblurring on the cryo-EM density map with a strength proportional to the pre-computed local resolution in each region of the map. LocSpiral [Kaur et al., 2021] employs the concept of spiral phase transformation to factorize the volume and enhance high-resolution features locally while preventing map distortions. LocScale [Jakobi et al., 2017] uses a sliding window approach to locally scale up the amplitudes of the density map to match with a pre-computed atomic reference structure. Although these local map sharpening techniques have achieved some success, they still have some drawbacks. LocScale requires the prior availability of atomic models, a condition that cannot be met most time. LocSpiral and LocalDeblur require a mask to differentiate the noise and macromolecules in the cryo-EM map, which is often not readily available.

To overcome these limitations, deep learning based approaches have been proposed for fully automated cryo-EM map sharpening. DeepEMhancer[Sanchez-Garcia et al., 2021] aims to mimic the local map sharpening effect of LocScale by training their U-Net [Ronneberger et al., 2015] based deep neural network using LocScale generated maps. Since the target LocScale generated maps were created from experimental maps, lower quality experimental maps can limit DeepEMhancer’s performance. Therefore, although DeepEMhancer performs well on some experimental cryo-EM maps, it faces difficulty in robustly improving a wide range of cryo-EM maps. In contrast, EMReady [He et al., 2023] uses a Swin-Conv-Unet [Zhang et al., 2023] based deep neural network trained on simulated maps that were generated from known protein structures using a reference Gaussian function. EMReady trained on the simulated maps can robustly improve the quality and interpretability of a wide range of experimental cryo-EM maps.

Following the approach of EMReady using simulated maps generated from known protein structures as labels to train deep learning methods, we develop CryoTEN - a new three-dimensional U-Net style Transformer equipped with Efficient Paired Attention (EPA) [Shaker et al., 2022] for enhancing cryo-EM density maps. CryoTEN robustly improves the interpretability of experimental cryo-EM maps. The EPA attention helps in effectively learning both spatial and channel-wise discriminative features. U-Net style skip connections aid in retaining the spatial information between the downsampling and upsampling layers and thus enhance the cryo-EM map without significant loss of information during the encoding and decoding process. CryoTEN was evaluated on a diverse set of experimental cryo-EM primary and half-maps using various map-model validation metrics. The results show that the quality of the maps processed by CryoTEN is substantially better than the original experimental cryo-EM maps and can be used to build better structural models than the original ones.

## Materials and Methods

### Data Collection

The advanced search tool of the RCSB Protein Data Bank (PDB) was used to filter PDB structures built from deposited single-particle cryo-EM maps that have a resolution between 2Å and 7Å. For PDB structures with multiple associated cryo-EM maps, only one cryo-EM map was chosen. Similarly, for cryo-EM maps with multiple associated PDB structures, only one structure was chosen. PDB structures with non-orthogonal map axes were removed. Protein sequences in the FASTA format and deposited cryo-EM primary maps were fetched for the filtered PDB structures from the PDB and Electron Microscopy Data Bank (EMDB) respectively. We used phenix.map model cc [Afonine et al., 2018] to compute Cross Correlation (CC) scores between PDB structures and associated deposited cryo-EM maps. To ensure the quality of the data, only maps that have CC mask value *>* 0.7 and CC box value *>* 0.6 were selected. Finally, to remove redundant cryo-EM maps that have similar sequences/structures, we used MMseqs2 to cluster their respective PDB structures that have a sequence identity *>* 30%, and only one structure was selected per cluster. The final non-redundant data collection consists of 1,521 PDB structure and map pairs, of which we randomly selected 1,295 maps as the training set, 76 maps as the validation set, and 150 maps as the test set.

### Data Processing

To train CryoTEN, the deposited experimental cryo-EM primary maps were used as input, and high-quality simulated maps generated from the corresponding PDB structures were used as targets (labels). These target simulated maps (label) were computed from PDB structures using a reference Gaussian function [DiMaio et al., 2009] as follows:

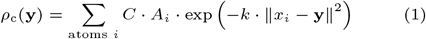

The density value *ρ*_c_(**y**) for a grid point with coordinate **y** in the simulated map is computed according to its distance with every atom in the PDB structure, where *x*_*i*_ is the coordinates of the *i*^*th*^ atom in the PDB structure and *A*_*i*_ is the atomic number of the *i*^*th*^ atom; *k* = (*π/*(0.9*R*_0_)^2^, where *R*_0_ is the unmasked Fourier Shell Correlation resolution at 0.143 threshold (FSC@0.143) of the deposited cryo-EM primary map computed using phenix.mtriage [Afonine et al., 2018] tool; and *C* = (*k/π*)^3*/*2^. The density value *ρ*_c_ is computed for all grid points to construct the simulated map.

Both the deposited cryo-EM primary maps and simulated density maps were resampled to a common, standardized grid size of 1Å. Density values of deposited cryo-EM maps were normalized to a range of 0 to 1 by the 99.999^th^percentile density value. Since cryo-EM maps are different in size and often large, we split all the maps into smaller blocks (cubes) so that each of them does not require much GPU memory to hold. Specifically, in the training set, deposited cryo-EM primary maps and simulated maps were initially split into overlapping blocks of size 64 x 64 x 64 with a stride length of 50, and were randomly cropped to blocks of size 48 x 48 x 48 on the fly during training to reduce overfitting. Only the blocks that contain protein structures were selected for training, while empty blocks were ignored. In the validation and test set, the maps were directly split into overlapping blocks of fixed size 48 x 48 x 48 with a stride length of 38. An overview of the data preparation process is illustrated in Figure 1a.

**Fig 1.**
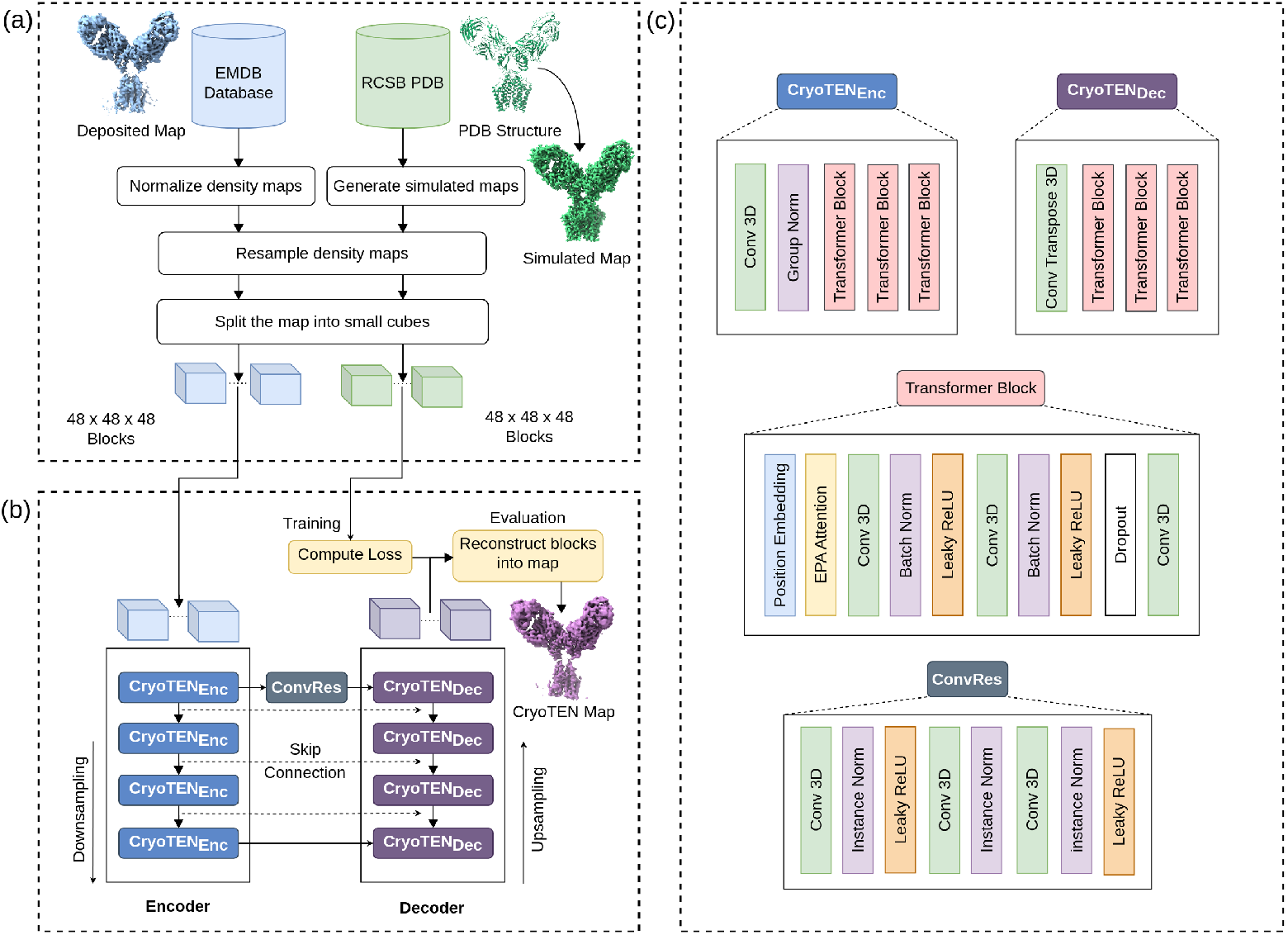
Overview of data processing and CryoTEN model architecture. **(a)** Data collection and preprocessing. **(b)** The CryoTEN model architecture along with the training and evaluation pipeline. **(c)** The structure of the CryoTEN model’s encoder, decoder, and residual convolution (ConvRes) block.

### Neural Network Architecture

CryoTEN is a U-Net style transformer-based deep neural network. It consists of four pairs of transformer-based encoders and decoders with U-Net like skip connections between the encoders and decoders as shown in Figure 1b. An encoder consists of a down-sampling convolution block, batch normalization, and three transformer blocks. A decoder consists of an upsampling convolution transpose block and three transformer blocks. Transformer blocks are equipped with positional embedding, convolution block, Leaky ReLUs (Rectified Linear Unit), and Efficient Paired Attention (EPA)[Shaker et al., 2022] along with batch normalization and dropouts for regularization as shown in Figure 1c. The first encoder and the last decoder are connected through an intermediate bottleneck block made up of a residual convolution block called ConvRes (see its structure in Figure 1c). ConvRes block comprises 3 pairs of convolution and instance normalization layers, two Leaky ReLU activation layers. In the ConvRes block, there is also a residual connection between the input and the third pair of convolution and instance normalization layers.

### Training and Validation

To reduce over-fitting during training, we augmented the training data by randomly cropping 48 x 48 x 48 density map blocks from 64 x 64 x 64 density map blocks. We also applied random 90-degree rotation and random axis flipping to the training data to further augment it. During training, the blocks of the experimental cryo-EM primary maps were used as input to CryoTEN. The output generated by CryoTEN was compared against the corresponding simulated density blocks (labels) to compute the loss. The loss is calculated as the masked mean square loss, where voxels being zero in both output blocks and simulated density blocks are masked and the loss for the remaining voxels after masking is calculated using the following equation:

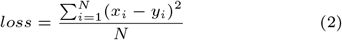

where *N* is the total number of voxels after masking, *x*_*i*_ is the value of the *i*^*th*^ voxel in the output map block and *y*_*i*_ is the value of the *i*^*th*^ voxel in the simulated density block. The Adam optimizer with an initial learning rate of 0.0005 was used to train the model to minimize the loss. To avoid over-fitting, the model was validated on the validation set every 5 epochs, and the learning rate was halved till 0.00001 if the validation loss did not improve in two consecutive times. CryoTEN was trained for 244 epochs on 4x NVIDIA A10 with a batch size of 15.

## Results

After training and validation, we blindly evaluated CryoTEN on the independent test set. During the evaluation, each input cryo-EM map was sliced into 48 x 48 x 48 blocks, which were fed as input to CryoTEN in batches. The output blocks generated by CryoTEN were then assembled to reconstruct the enhanced full density map of the same size as the input density map. The quality of the enhanced maps was then assessed.

### Evaluation on CryoEM primary maps

CryoTEN is first evaluated on the test set containing 150 cryo-EM deposited primary maps using various map-model validation metrics. Fourier Shell Correlation (FSC) of CryoTEN enhanced maps and original deposited cryo-EM primary maps are computed by comparing them with their corresponding atomic protein structures using phenix.mtriage [Afonine et al., 2018] tool. The unmasked map-model FSC resolution at both 0.143 (FSC@0.143) and 0.5 (FSC@0.5) thresholds are reported. Similarly, three Cross Correlation (CC) scores: CC box, CC mask, and CC peaks are computed for the maps by comparing them against their corresponding atomic structure using phenix.map model cc [Afonine et al., 2018] tool. The phenix.map model cc tool creates a molecular mask around the macromolecule (the determined protein structure). It also builds a map from the macromolecule called the model-calculated map. CC box is the CC computed between the entire experimental map and the model-calculated map. CC mask is computed between the experimental and model-calculated maps for regions only inside the molecular mask. CC peaks is computed around the union of regions having the N highest density values in the model-calculated and experimental target maps, where N is the number of grid points present inside the molecular mask. Finally, to measure the resolvability of atoms in the maps, the Q-score [Pintilie et al., 2020] metrics is computed using UCSF Chimera [Pettersen et al., 2004] mapq plugin. The average scores of these metrics for the original deposited primary maps and the CryoTEN processed primary maps are reported in Table 1.

**Table 1.** The comparison of the average map-model validation metrics scores of the 150 deposited cryo-EM primary maps in the test dataset and the corresponding CryoTEN processed maps. The scores are computed by comparing the maps with their respective atomic structures retrieved from the PDB.

| Metrics | Deposited | CryoTEN |
| --- | --- | --- |
| FSC@0.143 (Å) | 3.55 | 2.45 |
| FSC@0.5 (Å) | 6.38 | 3.73 |
| CC_Box | 0.7231 | 0.8508 |
| CC_Mask | 0.7874 | 0.7685 |
| CC_Peaks | 0.6439 | 0.7475 |
| Q-score | 0.5378 | 0.5435 |

The average map-model FSC@0.143 resolution of the CryoTEN processed cryo-EM primary maps is 2.45Å, 30.99% better than 3.55Å of the deposited cryo-EM primary maps. Similarly, the average map-model FSC@0.5 resolution of the CryoTEN processed maps is 3.73Å, 41.54% higher than 6.38Å for the deposited cryo-EM primary maps. Out of 150 maps, 100% of the CryoTEN processed maps exhibit an improvement in terms of the map-model FSC@0.143 resolution, and 96% of them show an improvement in terms of the map-model FSC@0.5 resolution.

In terms of the CC scores, the CryoTEN processed maps have substantially higher average CC box and CC peaks scores than the deposited primary maps, while their average CC mask score is slightly lower. CryoTEN achieves an average CC box score of 0.8508, 17.66% higher than 0.7231 of the deposited cryo-EM maps, indicating that it improves the overall quality of the entire maps. The average CC peaks score of CryoTEN processed maps is 0.7475, 16.09% higher than 0.6439 of the deposited cryo-EM maps, indicating that it improves the quality of the regions with the highest density that are important for structure determination.

However, the average CC Mask score of the CryoTEN processed maps is 0.7685, 2.4% lower than 0.7874 of the deposited cryo-EM maps, indicating that the cross correlation score over the region spanning the protein molecule is slightly reduced. But as the structure modeling results shown in Section “Improvement of protein structure modeling (map interpretability)” below, this minor reduction does not appear to affect the quality of the protein structural models built from the maps as the structures built from the CryoTEN-processed maps have better quality than the ones from the deposited maps.

Finally, the average Q-score of the CryoTEN processes maps is 0.5435, marginally better than 0.5378 of the deposited cryo-EM maps. Overall, the results from various map-model validation metrics above indicate that CryoTEN can reliably improve the quality of deposited cryo-EM primary maps. Figure 2 further compares the distribution of the map-model FSC resolution, Q-score, and CC scores between the deposited cryo-EM primary maps and CryoTEN processed maps.

**Fig 2.**
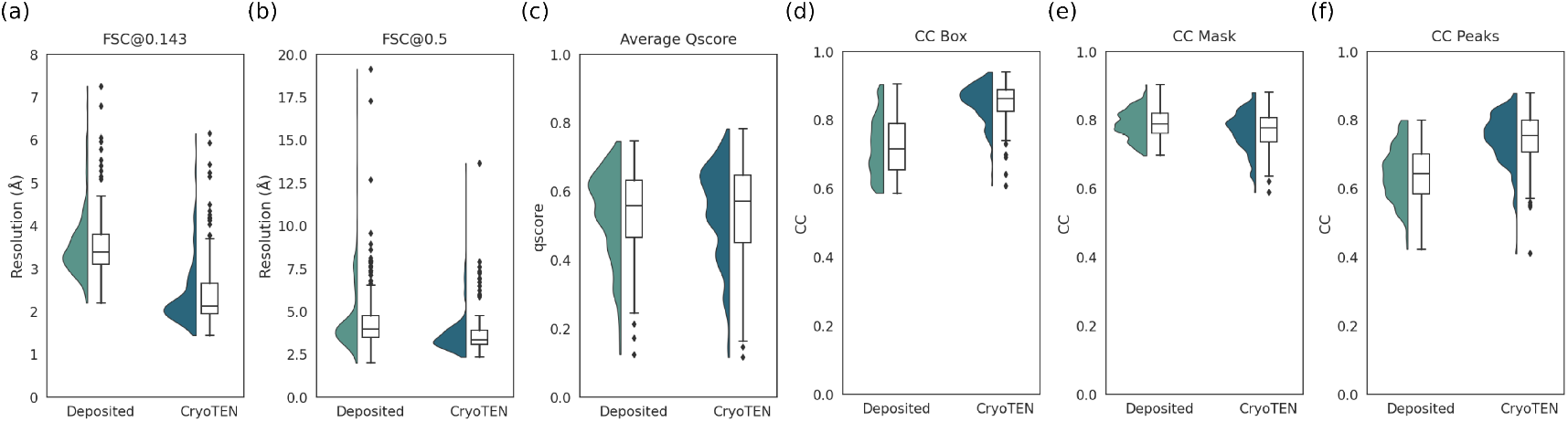
The comparison of the map-model validation metrics scores of the 150 deposited cryo-EM primary maps in the test dataset and the corresponding CryoTEN processed maps. Half violin and box plots of **(a)** unmasked FSC@0.143 resolution, **(b)** unmasked FSC@0.5 resolution (7 outliers with FSC@0.5 *>* 20Å in deposited primary maps are hidden for better visualization), **(c)** average Q-score, and **(d**,**e**,**f)** CC box, CC mask, CC peaks scores respectively.

### Evaluation on CryoEM half maps

During the 3D cryo-EM density map reconstruction process, in addition to constructing primary density maps from all 2D protein particle images, the protein particle images are usually split randomly into two subsets, each of which is used to build one separate 3D EM map called a half map in order to assess the reliability of the reconstruction according to the consistency between the two half maps. The unprocessed cryo-EM half maps tend to have lower resolution and higher noise than the primary maps because they have been post-processed using map sharpening techniques. Therefore, in addition to evaluating CryoTEN on experimental primary maps that have been processed by the Cryo-EM post-processing techniques, here we analyze how well CryoTEN can handle raw cryo-EM half maps that have not been processed by the cryo-EM post-processing techniques at all. Out of the 150 primary maps in the test set, 70 maps have corresponding unprocessed cryo-EM half map pairs available in EMDB. For each of them, one half map is chosen for the evaluation, resulting in 70 raw half maps for assessing CryoTEN. The results are reported in Table 2.

**Table 2.** Comparison of average map-model validation metrics scores of the 70 deposited cryo-EM half maps and CryoTEN processed half maps.

| Metrics | Deposited | CryoTEN |
| --- | --- | --- |
| FSC@0.143 (Å) | 3.75 | 2.61 |
| FSC@0.5 (Å) | 6.2 | 3.77 |
| CC_Box | 0.6796 | 0.8179 |
| CC_Mask | 0.7275 | 0.7152 |
| CC_Peaks | 0.5677 | 0.7038 |
| Q-score | 0.4651 | 0.4915 |

CryoTEN: Efficiently Enhancing Cryo-EM Density Maps using Transformers 5

Similar to the results on the primary density maps evaluation in previous section, the CryoTEN processed half maps have much higher average FSC@0.143 resolution, FSC@0.5 resolution, CC Box score, and CC Peaks score and moderately higher Q-score than the deposited half maps, while their average CC Mask score is slightly lower. The results show that, although CryoTEN is only trained on the deposited cryo-EM primary maps, it can also enhance unprocessed cryo-EM half maps. Since CryoTEN can enhance both processed and unprocessed cryo-EM maps, it can be used with existing post-processing tools such as RELION [Scheres, 2012, Rosenthal and Henderson, 2003] and CryoSPARC [Punjani et al., 2017] to improve the quality of any kind of density maps. Supplementary Figure 1 further compares the distributions of the FSC resolutions, Q-score, and CC scores of the deposited cryo-EM half maps and CryoTEN processed half maps.

### Improvement of protein structure modeling (map interpretability)

To evaluate the impact of CryoTEN on improving the quality of protein structures built from cryo-EM density maps (map interpretability), we performed chain-wise structural modeling on the 150 deposited cryo-EM primary maps in the test dataset and the corresponding CryTEN processed primary maps using an automatic de novo model building tool, phenix.map to model [Liebschner et al., 2019, Terwilliger et al., 2018a]. We used the zone tool in UCSF ChimeraX [Pettersen et al., 2021] to extract chain-wise map regions from the maps. The extracted chain-wise map regions and respective protein sequences of the chains were used by phenix.map to model to build the chain-wise atomic models. Out of all the chains extracted from 150 maps, phenix.map to model was able to build structural models for 700 chains from 124 maps successfully. To eliminate human influence, the modeled structures of the chains built from the 124 deposited maps and the corresponding CryoTEN processed maps are evaluated without any further model refinement.

The generated atomic model for each chain is evaluated using phenix.chain comparison tool [Liebschner et al., 2019] by comparing it against the respective structure of the chain from the PDB (considered as ground truth), and the average residue coverage score and sequence match score over all the chains are reported in Table 3. The residue coverage score (%) is the percentage of C*α* atoms in a modeled structure that are aligned within 3Å to its corresponding PDB structures. The sequence match score (%) is the percentage of residues in the modeled structure whose amino acid types exactly match those in the corresponding PDB structure.

**Table 3.** The quality (residue coverage and sequence match scores) of structural models of 700 protein chains built from the deposited 124 cryo-EM density maps and the CryoTEN processed maps.

| Maps | Residue Coverage(%) | Sequence Match(%) |
| --- | --- | --- |
| Deposited | 63.14 | 40.05 |
| CryoTEN | 71.79 | 43.52 |

The average residue coverage of the chain-wise atomic models built from the CryoTEN processed maps is 71.79%, 13.7% higher than 63.14% for the deposited cryo-EM maps, indicating that CryoTEN processing helps determine the positions of more C*α* atoms correctly. Moreover, the average sequence match score of the atomic models built from the CryoTEN processed maps is 43.52%, higher than 40.05% for the deposited primary maps, indicating that CryoTEN processing also improves the precision of determining the amino acid types of C*α* atoms.

For all 700 chains, CryoTEN processing improves the residue coverage for 84.86% of them, sequence match score for 60.14% of them, and either of the two for 91% of them. The results demonstrate that CryoTEN can reliably improve the quality of cryo-EM density maps for generating better structural models. However, it should be noted that here we only perform automatic de novo structure modeling to analyze the effects of CryoTEN enhancement of cryo-EM density maps. Further refinement of the structural models by human experts still plays a crucial role in building high-quality atomic models as fully automated density map-based structure modeling methods that match the quality of expert-built models are still not available despite the significant progress [Giri and Cheng, 2024b] in the area.

Figure 3 shows an example of CryoTEN substantially improving the deposited cryo-EM map (EMD-22338) of Epstein-Barr virus-encoded G protein-coupled receptor BILF1 at various contour levels by removing noise and adding structural details. The deposited maps and the CryoTEN enhanced maps are overlaid with their corresponding protein structure (PDB ID: 7JHJ) to visualize their quality. The regions where CryoTEN makes substantial improvement are circled. For instance, at a higher contour level, the circled high-density regions in the CryoTEN enhance map match the protein structure better than the deposited map in which some structural details are missing, while at a lower contour level, CryoTEN effectively reduces the background noise in the circle region. A similar example (EMD-22937 of H1 A/Michigan/45/2015 ectodomain) is illustrated in Supplementary Figure 2. The examples clearly illustrate how CryoTEN enhanced density maps can aid model building tools such as phenix.map to model to build better protein structures. Specifically, in the case of EMD-22338, the structural model of Chain R built from the CryoTEN enhanced map has a residue coverage of 78.7%, higher than 72.9% from the deposited Cryo-EM map. Moreover, the residue coverage of Chain A and D for the former is 75.2% and 85.4% respectively, substantially higher than 63.3%, and 37.3% for the latter.

**Fig 3.**
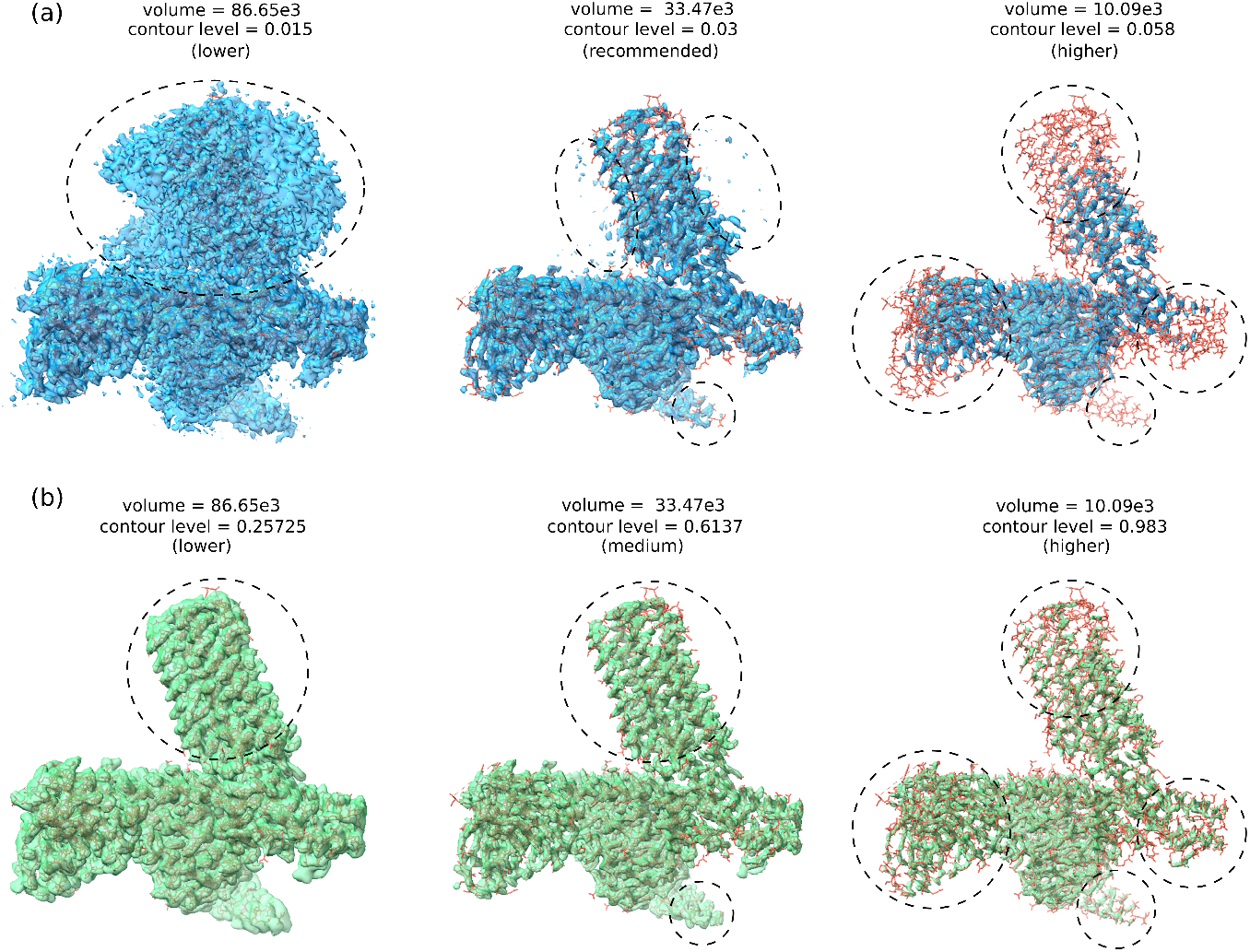
Comparison of an original deposited density map (EMD-22338) and its CryoTEN enhanced counterpart in superimposition with the known protein structure from the PDB. (a) The deposited density map (blue) is visualized at three contour levels (lower, recommended, and higher) in superimposition with the protein structure (PDB ID: 7JHJ). (b) The CryoTEN enhanced map (green) is visualized at three contour levels in superimposition with the protein structure. For direct comparison, the density volume of the two kinds of maps is set to equal at each contour level by adjusting the contour level of the CryoTEN enhanced map. The adjustment of the contour level is needed because the density values in the deposited map and the CryoTEN enhance map are not in the exact same range. The dotted circles in the images of the maps highlight the regions where CryoTEN substantially removes noise and/or adds more structural details. For instance, at the lower contour level, CryoTEN removes a lot of noise in the circled region, while at the high contour level, it adds some structural details that are missing in the circled regions of the deposited density maps

### Validation of robustness of CryoTEN in improving structure modeling (map interpretability)

To evaluate CryoTEN’s ability to enhance cryo-EM maps with varying FSC resolutions for structure modeling, we split the structural models built for the 700 chains from the 124 cryo-EM maps into three groups according to the resolution types of the original deposited density maps: low (4.5Å to 7Å), medium (3Å to 4.5Å), and high resolution (2Å to 3Å). The average residue coverage and sequence match scores for the structural models in each group built from the deposited maps and the CryoTEN enhanced maps are reported in Table 4. The results show that CryoTEN improves the quality of structural models in all three resolution groups. The improvement for the medium resolution group is most pronounced, where the residue coverage of models is improved by 9.76 percentage points and sequence match improved by 4.03 percentage points. Due to the poor resolvability of the atoms in low resolution maps, there is only a marginal improvement of 0.49 percentage points in the average sequence match score, but the residue coverage score is improved by 6.22 percentage points. Due to better atom resolvability in high resolution maps, the models built from experimental deposited maps achieve a good residue coverage and sequence match of 78.98% and 66.15% respectively, which CryoTEN was able to further improve to 84.28% and 68.37%, respectively. The results demonstrate that CryoTEN can robustly improve the quality of cryo-EM density maps to build better structural models.

**Table 4.** Robustness in improving map interpretability (structure modeling) for maps of various resolutions by CryoTEN. The 124 cryo-EM maps are grouped into three categories: low, medium, and high resolution, containing 10, 93, and 21 maps respectively. The average quality scores of the chain structural models built from the deposited cryo-EM maps and their CryoTEN enhanced counterparts in each category are shown.

| Map Resolution | Method | Residue Coverage(%) | Sequence Match(%) |
| --- | --- | --- | --- |
| Low<br>(4.5Å to 7Å) | Deposited | 27.99 | 7.03 |
|  | CryoTEN | 34.21 | 7.52 |
| Medium<br>(3Å to 4.5Å) | Deposited | 61.31 | 35.22 |
|  | CryoTEN | 71.07 | 39.25 |
| High<br>(2Å to 3Å) | Deposited | 78.98 | 66.15 |
|  | CryoTEN | 84.28 | 68.37 |

### Comparison with other deep learning methods

We compare CryoTEN with two existing deep learning methods for enhancing cryo-EM density maps, DeepEMhancer [Sanchez-Garcia et al., 2021] and EMReady [He et al., 2023] on the test dataset in terms of both the quality of the density maps and the empirical time complexity and memory consumption of the three methods (Table 5). While CryoTEN and EMReady processed all 150 maps in our test set, DeepEMhancer was able to process only 131 maps without crashing. Therefore, to make a fair comparison, we compare all the methods on only 131 maps in the test set in terms of average FSC@0.143, FSC@0.5, CC box, CC mask, CC peaks and Q-score. We also assess their running time and memory requirement on a subset of 20 test maps.

**Table 5.** Comparison of map-model validation metrics scores of 131 deposited cryo-EM primary maps in the test dataset, DeepEMhancer enhanced maps, EMReady enhanced maps, and CryoTEN enhanced maps as well as the processing time and GPU memory consumption of the three deep learning methods. The average processing time and GPU memory consumption for all three methods is benchmarked on the same subset of 20 maps with a batch size of 40.

| Metrics | Deposited | DeepEMHancer | EMReady | CryoTEN |
| --- | --- | --- | --- | --- |
| FSC@0.143 | 3.57 | 3.52 | 1.95 | 2.44 |
| FSC@0.5 | 5.81 | 4.71 | 3.17 | 3.77 |
| CC_Box | 0.7224 | 0.6309 | 0.8619 | 0.8499 |
| CC_Mask | 0.784 | 0.6096 | 0.7913 | 0.7669 |
| CC_Peaks | 0.6381 | 0.5733 | 0.7683 | 0.7433 |
| Q-score | 0.5313 | 0.4248 | 0.5867 | 0.5365 |
| Avg Processing Time (mins) | - | 43.27 ± 9.41 | 19.65 ± 17.95 | 1.66 ± 0.42 |
| GPU Memory Consumption (GB) | - | 21.88 | 8.86 | 3.41 |

The results show that EMReady improves the quality of the deposited density maps in terms of all the metrics, CryoTEN in terms of all the metrics except CC mask, and DeepEMhancer in terms of resolution (FSC@0.143 and FSC@0.5) but not cross correlation metrics (CC box, CC mask, CC peaks) and Q-score. CryoTEN performs substantially better than DeepEMHancer in terms all the metrics. For instance, the FSC@0.143 and CC Box of CryoTEN is 2.44Å and 0.8499, substantially higher than 3.52Å and 0.6309 of DeepEMhancer. The performance of CryoTEN is relatively close to the best-performing EMReady in terms of all the metrics, and both of them substantially increase FSC@0.143, FSC@0.5, CC box, and CC peaks of the maps. EMReady moderately increases Q-score, while CryoTEN only slightly improves it. EMReady has a CC mask score of 0.7913, sightly higher than 0.784 of the deposited maps, while CryoTEN has a slightly lower CC mask score of 0.7669. Because the quality of the density maps is indeed substantially improved by both EMReady and CryoTEN in terms of most of the metrics and better protein structures can be built from the CryoTEN processed maps, it is reasonable to hypothesize that CC mask is not sensitive to the change of the quality of density maps.

The execution time and memory consumption of the three deep learning methods were empirically compared on a subset of 20 density maps in the test dataset. CryoTEN runs much faster and requires much less GPU memory than EMReady and DeepEMhancer. On average, CryoTEN takes only 1.66 minutes to enhance a map with a standard deviation of ±0.42 minutes, whereas EMReady takes 19.65 minutes per map with a standard deviation of ±17.95 minutes and DeepEMhancer takes 43.27 minutes per map with a standard deviation of ±9.41 minutes. Moreover, on an NVIDIA A10 GPU, with batch size configured as 40 for all methods, CryoTEN consumes only 3.41 GB of GPU memory, compared to 21.88 GB of DeepEMhancer and 8.86 GB of EMReady. The per-map runtime and memory consumption for DeepEMhancer, and EMReady are reported in Supplementary Table 1. Therefore, CryoTEN is suitable for high-throughput enhancement of cryo-EM density maps where the processing speed is critical.

## Conclusion

In this study, we introduce CryoTEN, a U-Net style transformer equipped with the EPA attention mechanism to enhance cryo-EM density maps. CryoTEN is trained on a large non-redundant diverse set of 1,295 deposited cryo-EM primary maps and extensively evaluated on an independent test set. It can improve the quality of both primary density maps and half density maps in terms of multiple map-model validation metrics. The automatic de novo structure modeling experiment shows that the CryoTEN processed maps can be used to build better protein structural models than the original density maps. Moreover, CryoTEN runs more than 10 times faster and requires much less GPU memory than the existing deep learning methods, while still achieving a rather good performance in enhancing the quality of cryo-EM density maps.

## Data availability

The fine-tuned model weights are available at https://zenodo.org/records/12693785/files/cryoten.ckpt. The dataset used for training and evaluating CryoTEN can be downloaded using the scripts provided in our GitHub repository: https://github.com/jianlin-cheng/cryoten.

## Author contributions statement

J.C. conceived the project. J.S. collected the data and carried out the experiments under the guidance of J.C. and L.W. J.S., J.C. and L.W. wrote the manuscript. All authors reviewed the manuscript.

## Acknowledgments

This work is supported by an National Institutes of Health (NIH) grant (grant number: R01GM146340).

