## Supplementary Information for "CryoTEN: Efficiently Enhancing Cryo-EM Density Maps Using Transformers"

### Supplementary Figure 1: Plots of map-model validation metrics computed on deposited cryo-EM half maps and CryoTEN processed half maps

Computed various map validation metrics on deposited cryo-EM half maps (one of the half map pairs is used) and CryoTEN processed maps in our test set containing 70 half map pairs. Half violin and box plots of **(a)** unmasked FSC@0.143 resolution, **(b)** unmasked FSC@0.5 resolution (2 outliers with FSC@0.5 > 20Å in deposited half maps are hidden for better visualization), **(c)** average Q-score, **(d,e,f)** CC_box, CC_mask and CC_peaks scores respectively.

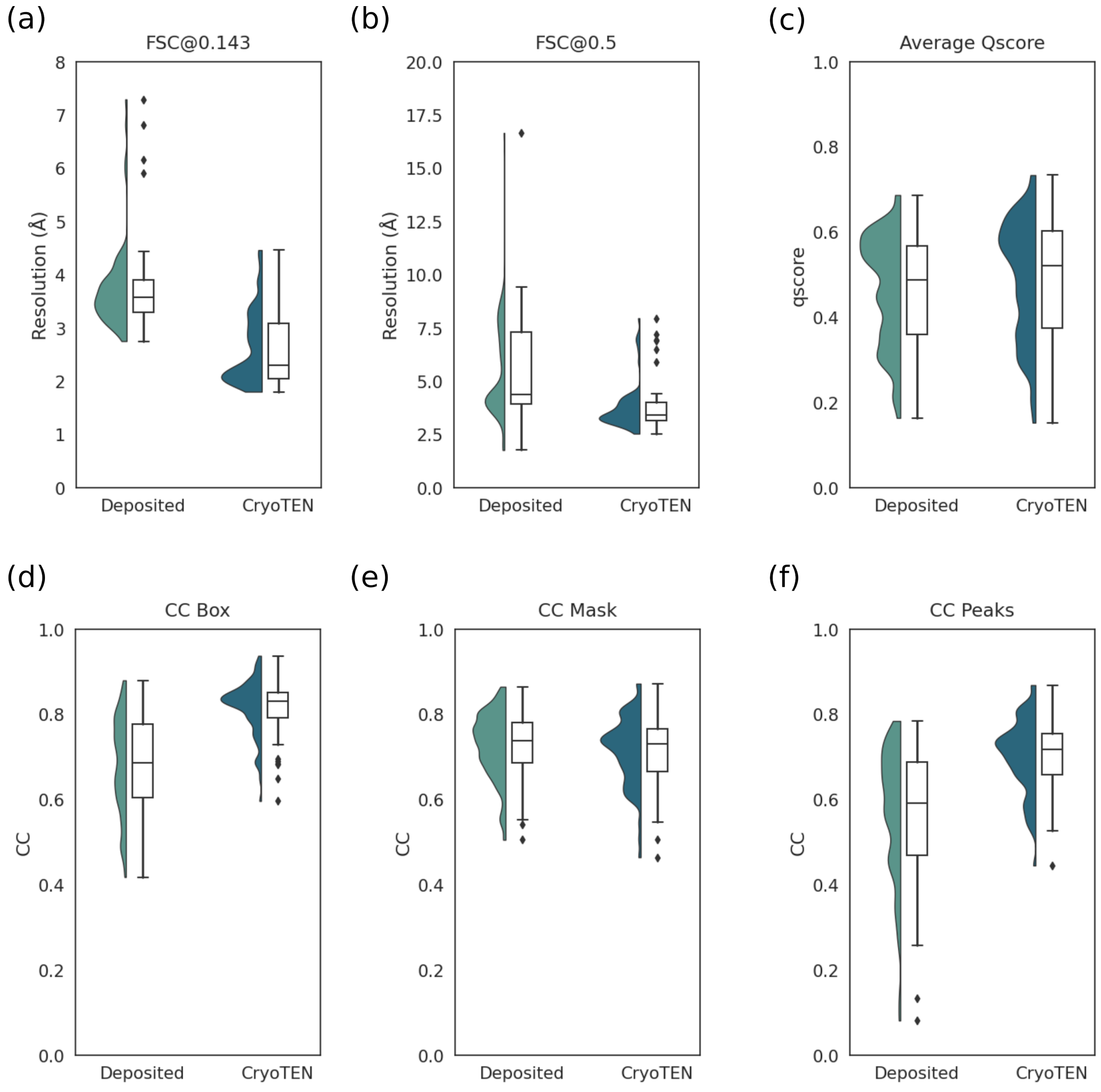

### Supplementary Figure 2: An example showing improvement in CryoTEN processed maps

An example that shows improvement in map quality when using CryoTEN. (a) and (b) compares the deposited density map (blue) and CryoTEN enhanced map (green) of EMD-22937 at various contour levels respectively. Each map overlapped with the corresponding PDB structure (PDB ID: 7KNA) (red). Since the density value distribution varies between maps, we match the density volume between the deposited and CryoTEN processed maps to make a fair comparison. The dotted circles in the images of the maps highlight the regions where CryoTEN substantially removes noise and/or adds more structural detail.

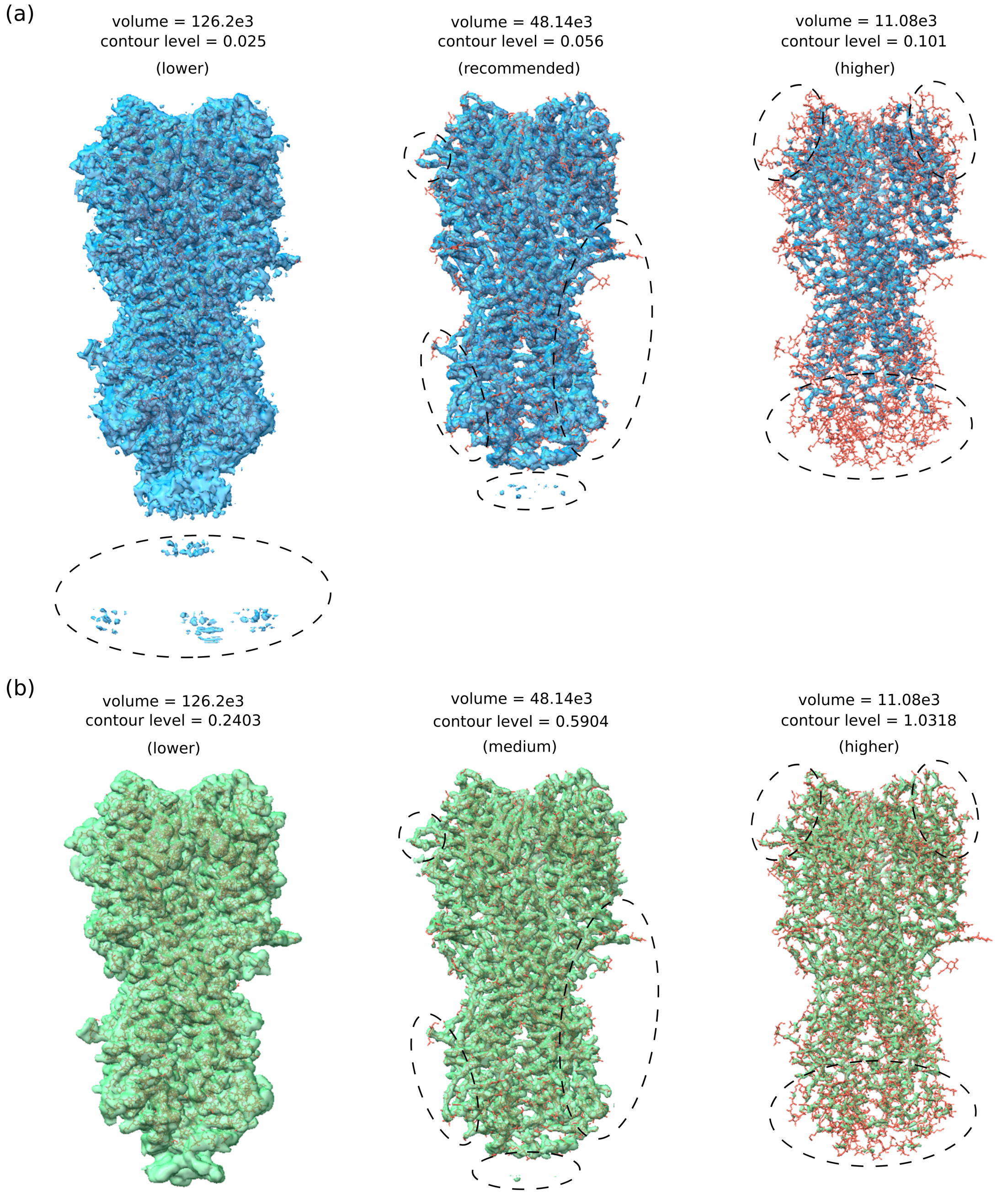

### Supplementary Table 1: Runtime Benchmark of CryoTEN, DeepEMHancer and EM-Ready on 20 maps

We ran CryoTEN, DeepEMhancer, and EMReady models on 20 maps in the test set with a batch size of 40. The time taken to process each map by these models is tabulated below. From the total time taken to process 20 maps, we can observe that CryoTEN performed significantly faster than DeepEMhancer and EMReady.

| EMDB ID | CryoTEN Time Taken (mins) | DeepEMhancer Time Taken (mins) | EMReady Time Taken (mins) |
|---|---|---|---|
| 35363 | 2.010357 | 54.836444 | 14.353148 |
| 26831 | 1.214203 | 37.408689 | 1.209833 |
| 27630 | 1.569806 | 38.069682 | 32.110613 |
| 23092 | 1.492781 | 48.146252 | 8.832931 |
| 13867 | 1.193822 | 41.560315 | 30.045555 |
| 23970 | 1.173997 | 38.321360 | 20.812409 |
| 26978 | 2.143782 | 37.432961 | 5.202853 |
| 22884 | 1.896650 | 54.818219 | 2.000178 |
| 32765 | 2.644996 | 47.408696 | 61.382500 |
| 24784 | 1.632092 | 42.320093 | 31.831395 |
| 32328 | 1.292059 | 40.041838 | 2.611483 |
| 4032 | 2.155701 | 70.056347 | 22.552169 |
| 26806 | 1.557495 | 45.533855 | 6.369265 |
| 20226 | 1.344403 | 35.729695 | 5.676419 |
| 33242 | 1.560156 | 34.966081 | 36.235096 |
| 14847 | 1.303207 | 47.767493 | 7.853368 |
| 34369 | 1.601426 | 35.538966 | 25.820971 |
| 10800 | 2.093365 | 50.292193 | 16.810930 |
| 15531 | 1.230027 | 28.490879 | 2.447058 |
| 32940 | 2.177127 | 36.571903 | 58.840337 |
| **Total (mins)** | 33.29 | 865.31 | 393.0 |
